# Targeted Clinical Metabolomics Platform for the Stratification of Diabetic Patients

**DOI:** 10.1101/664052

**Authors:** Linda Ahonen, Sirkku Jäntti, Tommi Suvitaival, Simone Theilade, Claudia Risz, Risto Kostiainen, Peter Rossing, Matej Orešič, Tuulia Hyötyläinen

## Abstract

**Background:** Several small molecule biomarkers have been reported in the literature for prediction and diagnosis of (pre)diabetes, its co-morbidities and complications. Here, we report the development and validation of a novel, quantitative, analytical method for use in the diabetes clinic. This method enables the determination of a selected panel of 36 metabolite biomarkers from human plasma.

**Methods:** Based on a review of the literature and our own data, we selected a panel of metabolites indicative of various clinically-relevant pathogenic stages of diabetes. We combined these candidate biomarkers into a single ultra-high-performance liquid chromatography-tandem mass spectrometry (UHPLC-MS/MS) method and optimized it, prioritizing simplicity of sample preparation and time needed for analysis, enabling high-throughput analysis in clinical laboratory settings.

**Results:** We validated the method in terms of limit of (a) detection (LOD), (b) limit of quantitation (LOQ), (c) linearity (*R^2^*), (d) linear range, and (e) intra- and inter-day repeatability of each metabolite. The method’s performance was demonstrated in the analysis of selected samples from a diabetes cohort study. Metabolite levels were associated with clinical measurements and kidney complications in type 1 diabetes (T1D) patients. Specifically, both amino acids and amino acid-related analytes were associated with macro-albuminuria. Additionally, specific bile acids were associated with kidney function, anti-hypertensive medication, statin medication and clinical lipid measurements.

**Conclusions:** The developed analytical method is suitable for robust determination of selected plasma metabolites in the diabetes clinic.

## Background

The incidence of type 2 diabetes (T2D) is rising globally, currently estimated to exceed 450 million patients worldwide. In addition, the prevalence of prediabetes is approximately 2-3 times higher than for diabetes. Prediabetes is a condition with a high risk of progression to T2D, with a yearly conversion rate of 5%-10% *(1, 2).* It is also known that excessive hepatic fat accumulation is a typical feature of T2D patients and plays an important, pathogenic role in disease development and progression. Particularly, non-alcoholic fatty liver disease (NAFLD) may have an important, deleterious impact on diabetic patients, increasing the risk of cardiovascular complications. Moreover, there is evidence of associations between prediabetes and complications of diabetes such as early nephropathy, small fiber neuropathy, early retinopathy and risk of macrovascular disease *(2)*. Therefore, there is a need for predictive tools for efficient and accurate tracking of the progression from the state of normal glucose tolerance (NGT) to pre-diabetes and finally to T2D, as well as a need for the identification of those individuals with T1D and T2D who are at risk of developing diabetic complications. There is also a need for improved stratification of those individuals who already have the disease based on their risk of developing complications. Finally, there is a pressing need to then tailor intervention strategies to these individuals. Ideally, knowledge about the underlying pathophysiological characteristics associated with either fasting or postprandial glucose dysregulation would be utilized in order to optimize the efficacy of any interventions *(3).*

The complex etiology of diabetes makes effective screening, diagnosis, prognosis and intervention challenging. Several studies have shown changes in the circulating levels of specific metabolites prior to an individual developing overt T2D. For example, the Framingham Offspring, European Investigation into Cancer and Nutrition (EPIC) Potsdam, Metabolic Syndrome in Men (METSIM), Cardiovascular Risk in Young Finns (CRY) and Southall and Brent Revisited (SABRE) studies have replicated the finding of increased levels of branched-chain amino acids and their derivatives, aromatic amino acids and α-hydroxybutyrate, even years ahead of conversion to overt T2D *(4–9).* Amino acids, particularly tyrosine, were found to be associated with risk of microvascular disease *(10).* Also other metabolites, *e.g.* 1,5-anhydroglucitol, norvaline and l-aspartic acid, were found to be associated with macroalbuminuric diabetic kidney disease *(11, 12),* while glutamine, glutamic acid and symmetric dimethylarginine (ADMA) were suggested as potentially-predictive biomarkers of diabetic complications *(13–15).* Several metabolites *(e.g.,* α-hydroxybutyrate, β-hydroxypyruvate and 1,5-anhydroglucitol (1,5-AG)), were associated with regulation of glycemic control *(16, 17).* Many lipids were identified as predictive biomarkers of diabetes. Specifically, triglycerides of low carbon number and double bond count as well as lysophosphatidylcholine, LPC(18:2), were identified as early predictors of T2D *(18, 19).* Notably, these markers were unaffected by obesity *(18).* Additionally, bile acids were associated with T2D and insulin resistance *(20, 21).* Mannose *(22),* 2-aminoadipic acid *(23, 24)* as well as indoxyl-sulfate and cresyl-sulfate *(26)* were suggested as possible biomarkers and, finally, creatinine(25) is already routinely implemented as an estimate of renal function.

Most of the studies described above have been performed with non-targeted metabolomics methods, using workflows which are difficult to apply in routine clinical laboratory settings. Herein, our goal was to develop a fast and robust method for quantitative analysis of a selected panel of metabolite biomarkers, which are informative as to the prediction and diagnosis of (pre)diabetes and its co-morbidities / complications, as well as in follow-up of interventions. We developed a method which includes 36 metabolites, representing several metabolite classes, including amino acids, bile acids, carnitines, phenolic compounds and small organic acids. The method is based on simple sample preparation and fast, quantitative ultra-high-performance liquid chromatography coupled to tandem mass spectrometry (UHPLC-MS/MS) analysis. Both sample preparation and the subsequent analyses were optimized and validated. Additionally, the method was demonstrated in a subset of samples from a cohort of diabetic patients, who were observed at the Steno Diabetes Center Copenhagen between 2009 and 2011 *(27).*

## Materials and methods

### Chemicals and standard solutions

Information on the chemicals used for this work is given in **Supplemental Methods**. Additionally, the structures of the compounds (**Supplemental Figures S1A-C**) and the preparation of standards and stock solutions is also described in **Supplemental Methods** and in **Tables 1** and **2**.

**Table 1.**
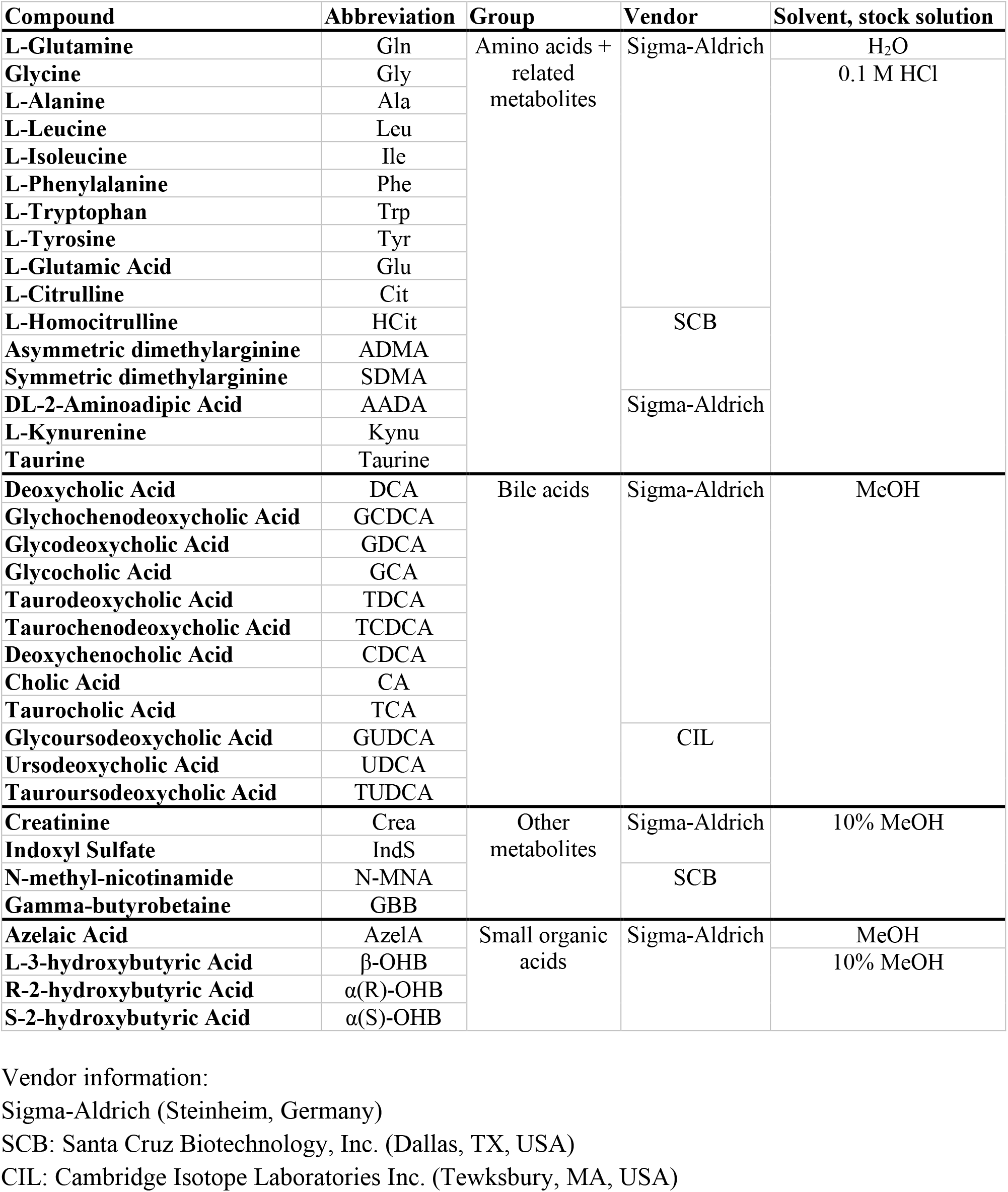
Standard compounds acquired for quality control and for quantitation.

**Table 2.**
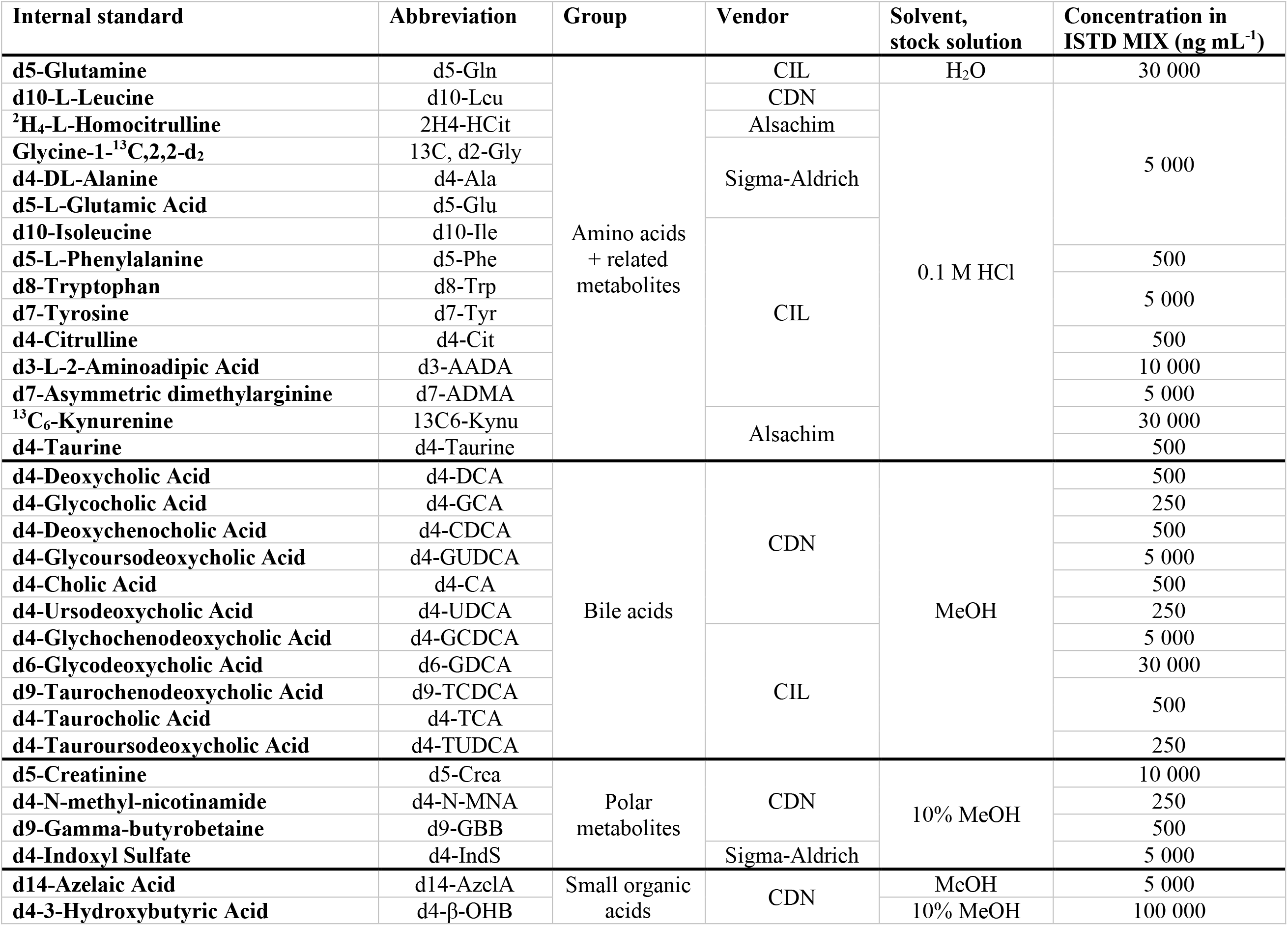
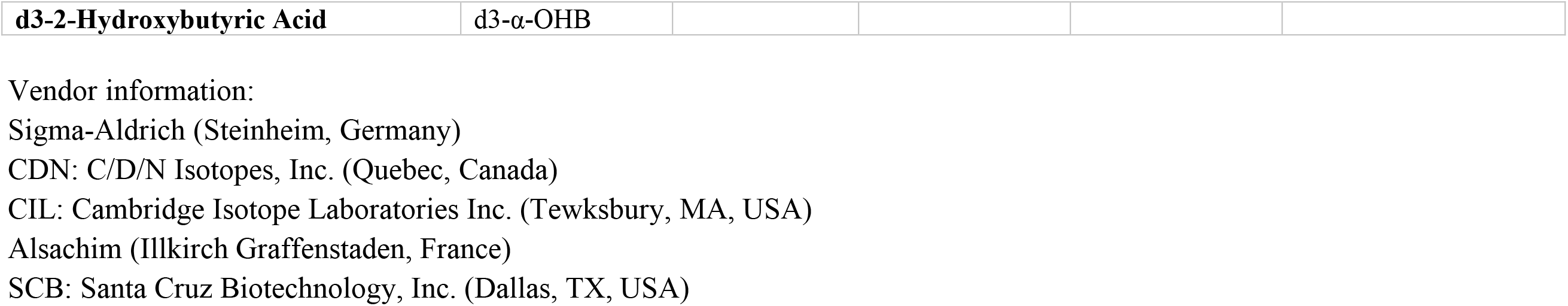
Internal standards, with concentrations in ISTD MIX, acquired for quality control and for quantitation.

### Samples

Plasma samples from a previously-described cohort *(27, 28)* were used for validation of the method. In short, during 2009-2011, a total of 1285 patients were invited to enter a study examining diabetic complications at the Steno Diabetes Center Copenhagen (SDCC). The study conformed to the Declaration of Helsinki and was approved by the Danish National Committee on Biomedical Research Ethics (2009-056; NCT01171248). Additionally, all patients gave written, informed consent. Of the invited 1285 patients, 676 accepted to participate and for our purposes, to demonstrate method functionality, a subset of 50 patient samples was analyzed. In addition to these plasma samples, pooled plasma samples from the SDCC were used for method development and validation as well as for quality control. All plasma samples were stored at −80 °C until analysis.

### Sample preparation

Sample preparation included protein precipitation and derivatization. 10 μL of 1 M 5-sulphosalisylic acid dehydrate (SSA) solution was added to 30 μL of plasma sample, samples were vortex mixed and centrifuged at 9000 RCF (5 minutes at 4 °C) after which 20 μL of the upper phase was collected, After addition of 20 μL of the ISTD MIX 20 μL of a 6-aminoquinoline-N-hydroxy-succinimidyl carbamate-reagent (AQC-reagent) (5 mg mL^-1^, at 55 °C) was added, and the samples were vortex mixed and stored at −80 °C until analysis.

The samples in the validation study were randomized before sample preparation and again before analysis. Calibration curves were created at the beginning and at the end of the sample analyses. Additionally, blank samples and pooled plasma samples were included in the analytical sequence for quality control purposes. Samples were injected three times, resulting in three technical replicate measurements for each of the 50 samples.

### Ultra high-performance liquid chromatography (UHPLC) – Mass spectrometry

The UHPLC-QQQ-MS system is described in detail in **Supplemental Methods**. Briefly, the UHPLC analysis was performed on a Kinetex^®^ F5 column (100 x 2.1 mm, particle size 1.7 μm) with a flow rate of 0.4 mL min^-1^ and an injection volume of 2 μL. H2O + 0.1% HCOOH (A) and ACN:IPA (2:1, v/v) + 0.1% HCOOH (B) were used as the mobile phases for gradient elution. Both positive and negative ion mode MS- and MS/MS-spectra (scan range *m/z* 40-600) were acquired (**Supplemental Tables S1** and **S2**).

## Results

Based on our earlier diabetes-related studies, as well as on the results published in the literature, we selected 36 specific metabolites for this study (**Table 1**) *(2, 4–7, 9, 10, 13, 16–20, 22, 23, 31–33)*. Our aim was to develop a robust and fast analytical assay in terms of both sample preparation and analysis, for quantitative determination of these selected metabolites. However, analyzing both highly polar and nonpolar metabolites in a single method is highly problematic. As some of the candidate biomarkers (e.g., very polar sugar derivatives and neutral lipids such as triacylglycerols) would have required a second sample preparation step and/or analytical method, these were excluded from the final method. The method was validated in terms of (a) limit of detection (LOD), (b) limit of quantitation (LOQ), (c) linearity (*R*^2^) and linear range and (d) intra- and inter-day repeatability of each analyte.

### Sample preparation

The main goal in the selection of conditions for sample preparation was the development of a workflow that is simple, robust and feasible to automate, while taking into consideration the LC-MS method as well. Here, we combined a simple protein precipitation with acid followed by derivatization of amino acids and structurally-related compounds. For the protein precipitation, acidic conditions were chosen, as protein precipitation with methanol or acetonitrile would have required evaporation of the solvent prior to derivatization and analysis. The amount of derivatization reagent, the amount and type of the solvent and buffer as well as the time for the derivatization reaction were optimized. Since the derivatization reagent has an impact on the MS detection, the conditions were optimized to decrease ion suppression as well as to improve the overall robustness of the method. Dry ACN was used for dissolving the AQC-reagent, as even trace amounts of water in the solvent can react with the reagent. The final sample preparation conditions included protein precipitation with SSA, followed by neutralization and pH adjustment using a mixture of carbonate buffer and NaOH) prior to the derivatization with AQC in anhydrous ACN. The MS spectra showed that only amino acids and related compounds with amino acid functionality (namely the amino acids, AADA, ADMA, SDMA, kynurenine and taurine) were derivatized and not any of the other targeted compounds.

### LC-MS

MS- and MS/MS-spectra were acquired for each of the analytes in order to select optimal precursor and product ions for selected reaction monitoring (SRM) analyses (**Supplemental Figures S2A-E**). Depending on the ionization properties of the different analytes, protonated ([M+H]+) or deprotonated ([M-H]^-^) molecules were chosen as precursor ions (**Supplemental Tables S1** and **S2**). MS/MS-spectra were acquired and the most selective and intense product ions were selected for SRM analyses. When possible, one ion transition was chosen for quantification and another ion transition was chosen as the qualifying ion transition to ensure correct measurements of the analytes. Finally, the analysis parameters (fragmentor voltage, collision energy, cell accelerator voltage) were optimized for each ion transition (**Supplemental Tables S1** and **S2**). All the derivatized amino acids and related compounds (**Figure 1**) produced the product ion [M-H-170]^-^. These were then selected for SRM analyses together with one other diagnostic product ion (where possible). Among the bile acids, CDCA and UDCA were not fragmented and, therefore, the only chosen product ions for these two analytes were their deprotonated molecules. For isomeric compounds (GCDCA, GDCA and GUDCA; TCDCA, TDCA and TUDCA) the MS and MS/MS-spectra are similar to the same three main product ions and their separation depends on chromatographic separation. In addition, TCA shows the same three main product ions as TCDCA, TDCA and TUDCA, but has different precursor ion.

**Figure 1.**
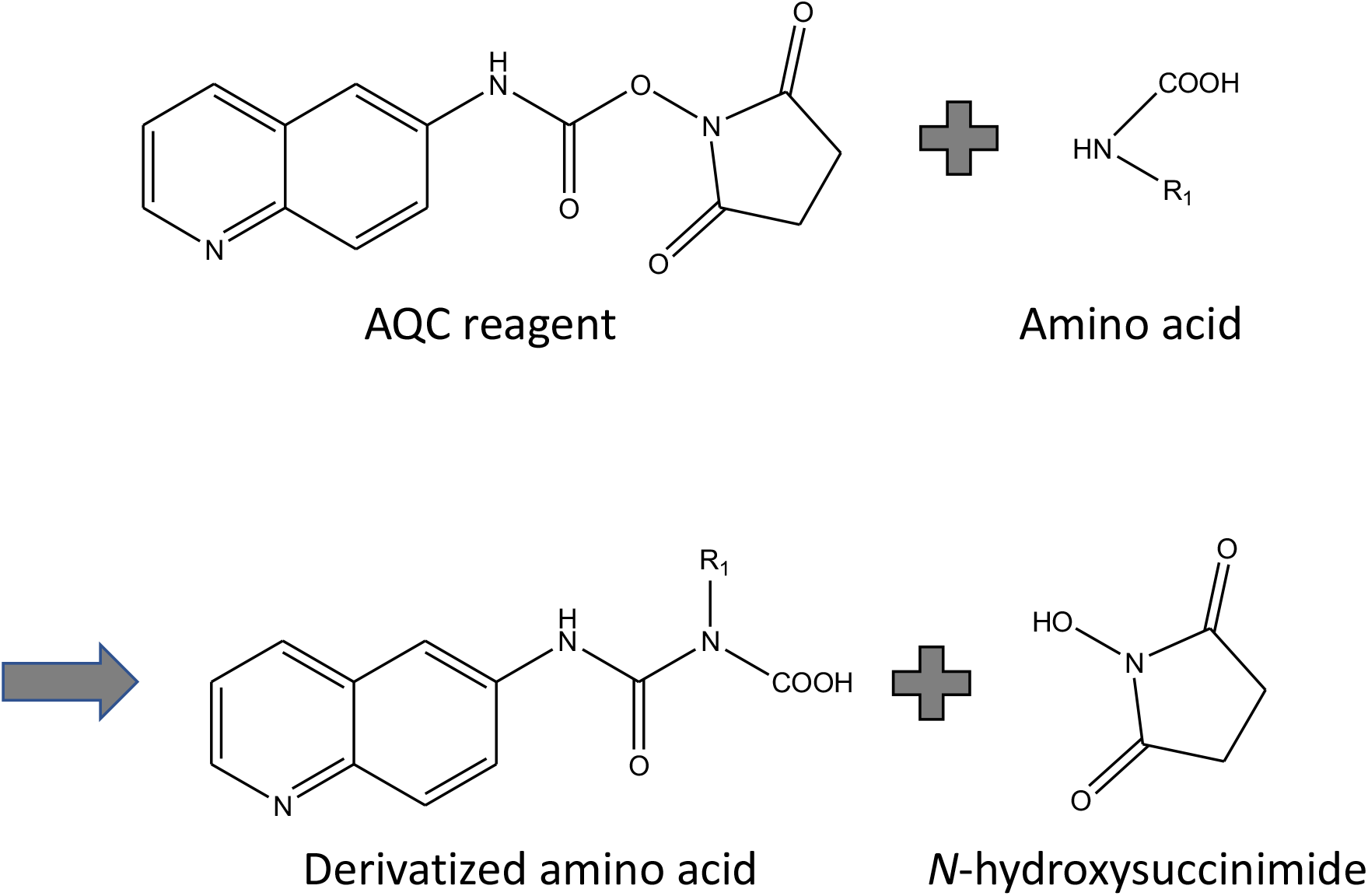
Derivatization reaction of amino acids and amino acid related compounds.

In the optimization of the LC-MS method, different columns (Ascentis Express RP-Amide, Poroshell 120 SB-AQ, Acclaim RSLC PolarAdvantage, Acclaim Trinity P2 and Kinetex^®^ F5 column) and different LC modes were tested. Based on the resolution of the chromatographic separation, the Kinetex^®^ F5 column was chosen for further optimization. The conditions were optimized to include sufficient retention for the most polar compounds. Therefore, the gradient elution was initiated at 99% of the aqueous eluent. The UHPLC method showed good chromatographic performance (**Figure 2**), fulfilling general acceptance criteria for an analytical method (**Section 3.3**). For a few of the analytes, the resolution was, however, insufficient to achieve baseline separation and due to very similar MS/MS-spectra these analytes (α(R)-OHB & α(S)-OHB, Leu & Ile, TDCA & TCDCA, GCDCA & GDCA, ADMA & SDMA) were quantified together.

**Figure 2.**
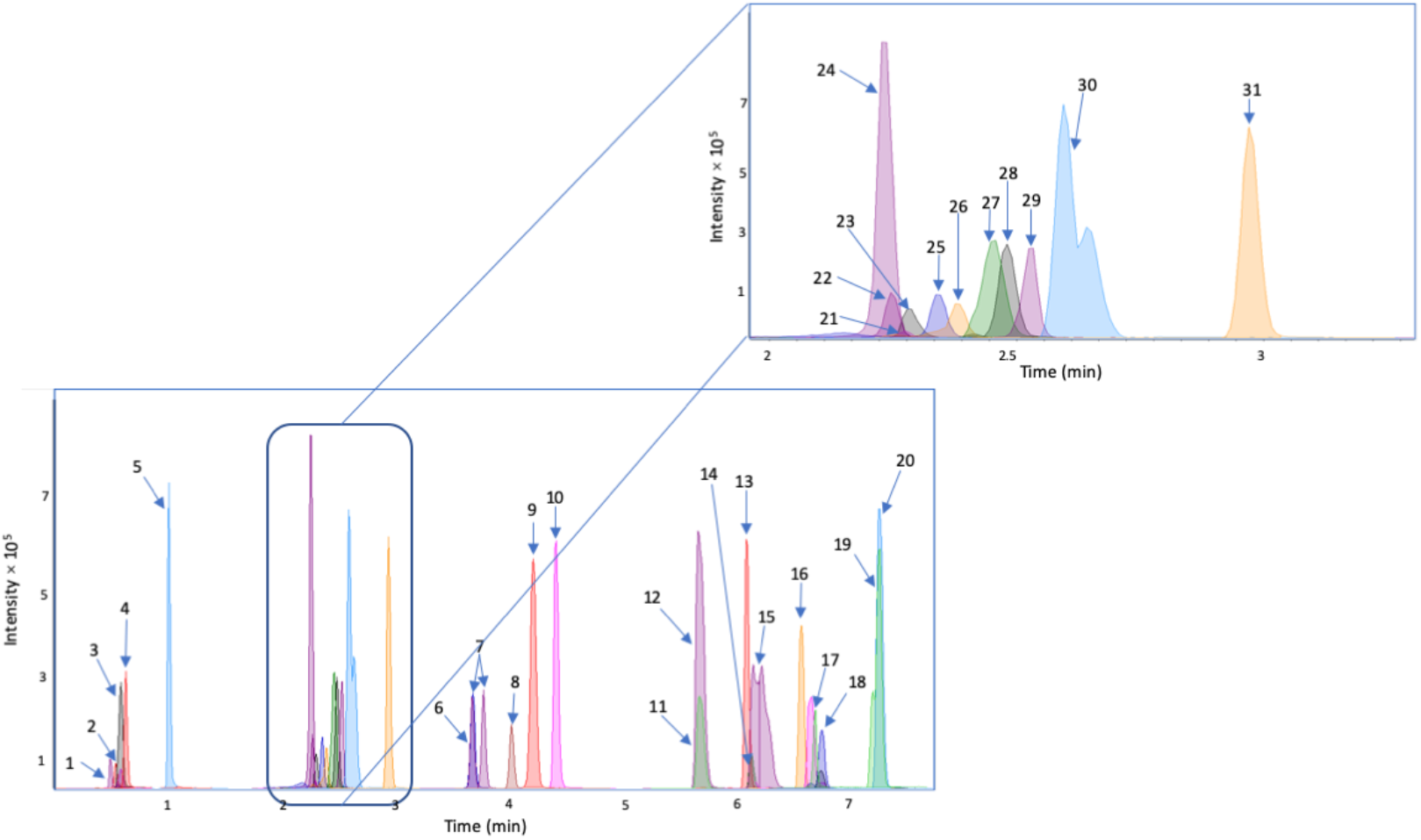
Chromatograms representing the chromatographic separation of the analytes. The peak numbers correspond to the following analytes: (1) Crea, (2) GBB, (3) α(R)-OHB & α(S)-OHB, (4) β-OHB, (5) N-MNA, (6) Kynu, (7) Leu & Ile, (8) Phe, (9) AzelA, (10) Trp, (11) TUDCA, (12) TCA, (13) GCA, (14) GUDCA, (15) TDCA & TCDCA, (16) CA, (17) CDCA, (18) GCDCA & GDCA, (19) UDCA, (20) DCA, (21) Gly, (22) Gln, (23) ADMA & SDMA, (24) Taurine, (25) Phe, (26) Gln, (27) HCit, (28) Ala, (29) AADA, (30) IndS and (31) Tyr.

### Method validation

The quantitative performance of the developed UHPLC-ESI-MS/MS method was evaluated with respect to (a) limit of detection (LOD), (b) limit of quantitation (LOQ), (c) linearity (*R*^2^) and linear range and (d) intra- and inter-day repeatability (**Table 3**). The LODs (at *S/N* ≥ 3) were measured from standard samples and are remarkably different for different analytes, with the lowest LOD being < 2.5 ng mL^-1^ (being the lowest measured concentration) for Ala, AzelA, GCDCA, GDCA, Leu, Ile, N-MNA, Phe and TCA. The highest LOD, on the other hand, was 25000 ng mL^-1^ for α(R)-OHB & α(S)-OHB. These results indicate an acceptable sensitivity within the concentration ranges normally detected in blood samples for most of the analytes, *(34)* except for α(R)-OHB & α(S)-OHB, for which the LODs are clearly higher than the lowest amounts previously detected in blood samples (3120 ng mL^-1^). Calibration curves and the intra- and inter-day repeatability were determined by using normalized peak areas. For the analytes which were quantified together *(i.e.,* GCDCA and GDCA, ADMA and SDMA, and TDCA and TCDCA), only one ISTD was used. The ISTDs used for GCDCA and GDCA, ADMA and SDMA, and TDCA and TCDCA were GDCA-d6, ADMA-d7 and TCDCA-d9, respectively. Additionally, for a few analytes (*i.e.,* GBB, Crea, α(R)-OHB, α(S)-OHB and β-OHB), the ISTD signal was not repeatable and therefore, the validation parameters of these analytes were measured without normalization to an ISTD. The calibration curves were determined within a concentration range of 2.5-75000 ng mL^-1^. The linear ranges showed a broad variation between the different analytes (**Table 3**). This variability is, however, acceptable since the biological concentration of each analyte is within the measured linear range. The coefficients of determination (*R*^2^) were higher than 0.97 for all analytes and above 0.99 for most analytes. This shows that the method has both good linearity and quantitation ability for each analyte, with accuracies well within the general requirement of 80-120%.

**Table 3.**
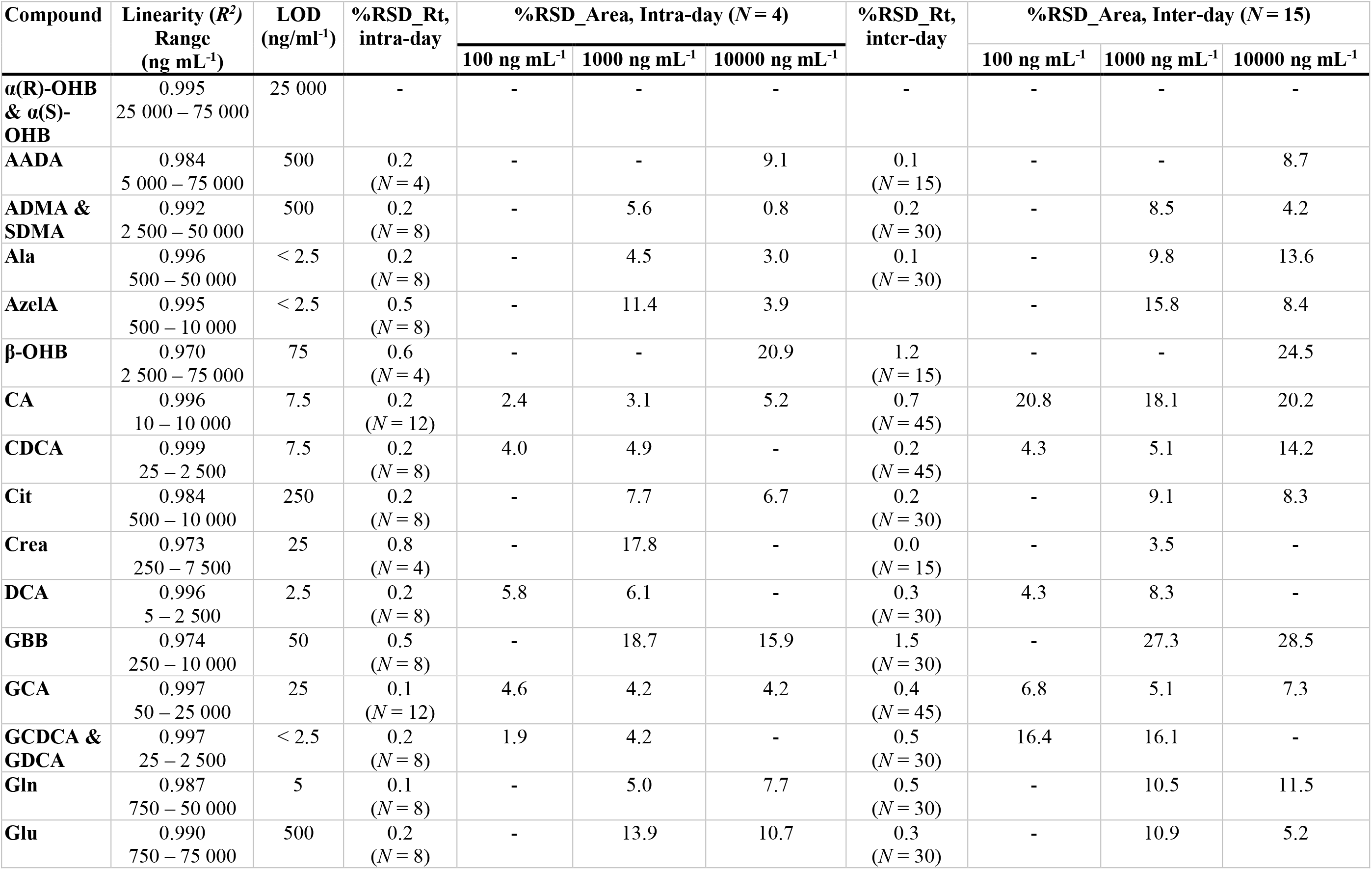
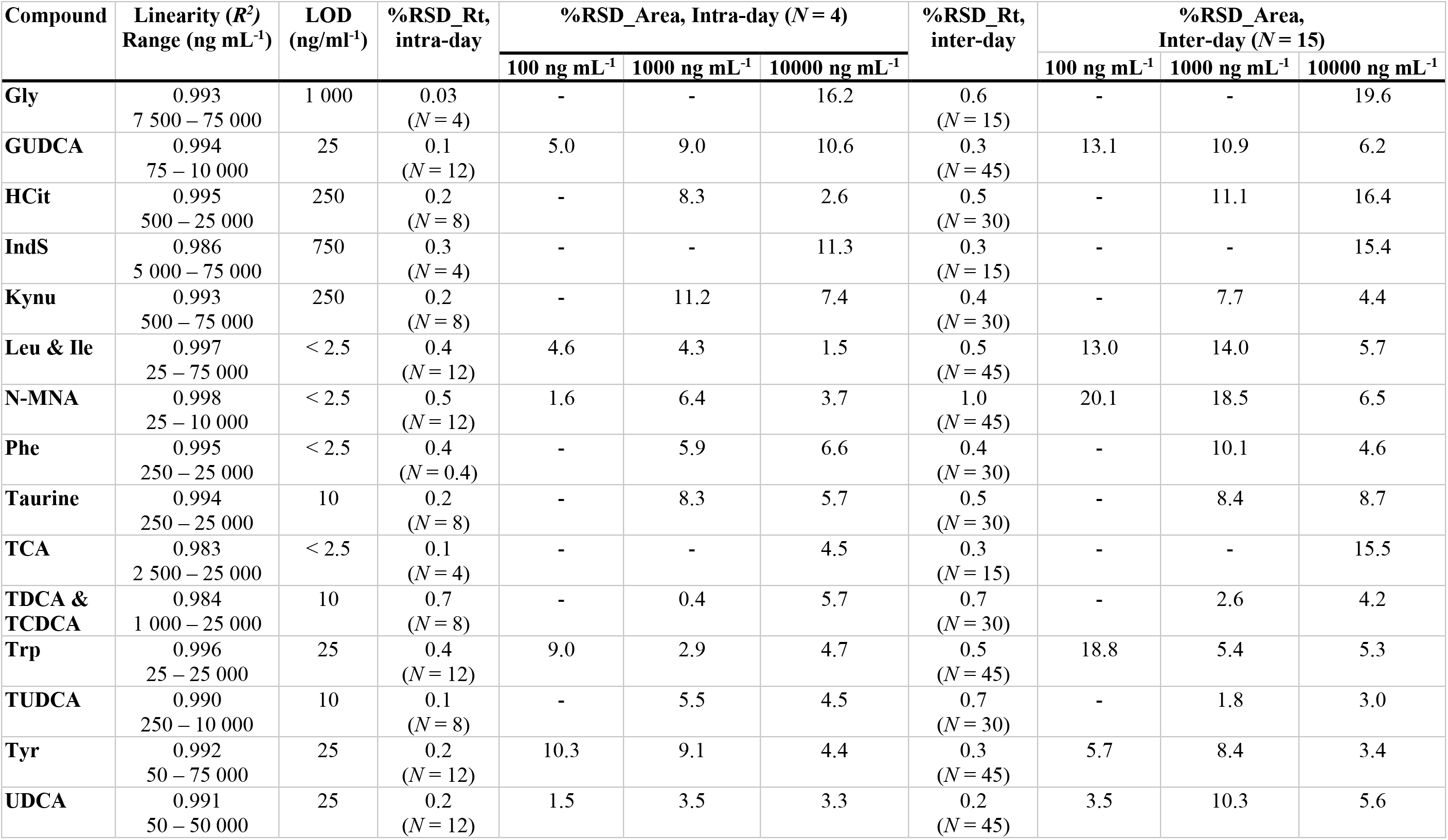
Limit of detection (LOD), limit of quantitation (LOQ), linearity (*R*^2^), linear range, intra- and inter-day repeatability.

For the repeatability studies, three standard samples (c= 100 ng mL^-1^, 1000 ng mL^-1^ and 10000 ng mL^-^)^1^ were analyzed in four consecutive runs and in three runs on five consecutive days for intraday and inter-day repeatability measurements, respectively. Relative standard deviations (%RSD) were calculated for both the intra- and inter-day studies (**Table 3**). The %RSDs for the intra-day repeatability studies were generally below 1.5% and 20.8% for the retention times and normalized peak area ratios, respectively. There are a few exceptions to these results for the analytes with no internal standards (Crea, GBB and β-OHB). The %RSDs for the intra-day and inter-day repeatability for these three analytes was between 17.8 and 20.9% and between 3.5 and 24.5%, respectively.

### Feasibility of the method for the analysis of samples from a diabetes cohort

The feasibility of the developed UHPLC-ESI-MS/MS method for the analysis of biological samples was demonstrated by analyzing plasma samples from individuals with diabetes who had a wide range of albuminuria. Albuminuria is a pathological condition where the protein albumin is present in the urine in abnormal amounts. It is a sign of diabetic kidney disease, which often occurs in especially subjects with type 1 diabetes *(28).* In healthy subjects (normo-albuminuric), only trace amounts of albumin (< 30 mg/24 hours) are present in the urine while subjects with elevated amounts of albumin in the urine, on the other hand, can be classified as either micro-albuminuric (c = 30 – 299 mg/24 hours) or macro-albuminuric (c ≥ 300 mg/24 hours).

In total, 50 samples were selected from a previously-described study cohort of a total of 676 participants *(28).* The subset was created with computational sampling, aiming at finding a small random subset of the cohort, where the distributions of potentially confounding clinical variables are as similar as possible between the two study groups. This allowed us to study associations between metabolites and albuminuria even in this small sample set whilst avoiding the confounding effects of other factors. The clinical variables assessed were age, antihypertensive medication, BMI, duration of diabetes, glycated hemoglobin (HbA1c), insulin day dose, sex, smoking, systolic blood pressure, total cholesterol and total triglycerides.

Selection of the best random subsample was done in four steps: (1) In total, 1 million n=25+25 sub-samples were drawn with random sampling, (2) the correlation between each clinical variable and the albuminuria group variable was computed for each subsample, (3) the highest absolute value of correlation in each subsample was identified, and (4) the random subsample with the lowest value of maximum correlation was selected for being the least-confounded random subset for analysis.

Computational selection resulted in a balanced subset of samples from 25 normo-albuminuric and 25 macro-albuminuric participants. The highest Pearson correlation to the albuminuria group variable among the clinical variables was 0.21 for total triglycerides. All other clinical variables had a lower absolute correlation to the group variable, suggesting that the selected small subset was not confounded by imbalance in the clinical characteristics.

Associations between metabolite concentrations and relevant clinical variables were tested with metabolite-specific mixed-effects models using the R-package limma *(35).* Metabolite concentrations entered the model as the dependent variable, participant identity as the random effect and the following clinical variables as fixed effects: albuminuria group, age, BMI, estimated globular filtration rate (eGFR; kidney function), glycated hemoglobin (HbA1c; glycemic control), sex, systolic blood pressure, total cholesterol, total triglycerides. Significance tests of coefficients were corrected for multiple testing over the metabolites with the Benjamini-Hochberg method *(36).*

Associations indicated by significant model coefficients (multiple-testing-corrected p < 0.05) were visualized as a bipartite network (**Figure 3**) between clinical variables and metabolites with the R-package ggplot2 *(37).* Strength (log-10-transformed coefficients) and the signs of each association were shown in the width and the color of the line, respectively. Metabolomic associations to albuminuria group and eGFR, which are the key variables of the present study, were highlighted with opaque lines.

**Figure 3.**
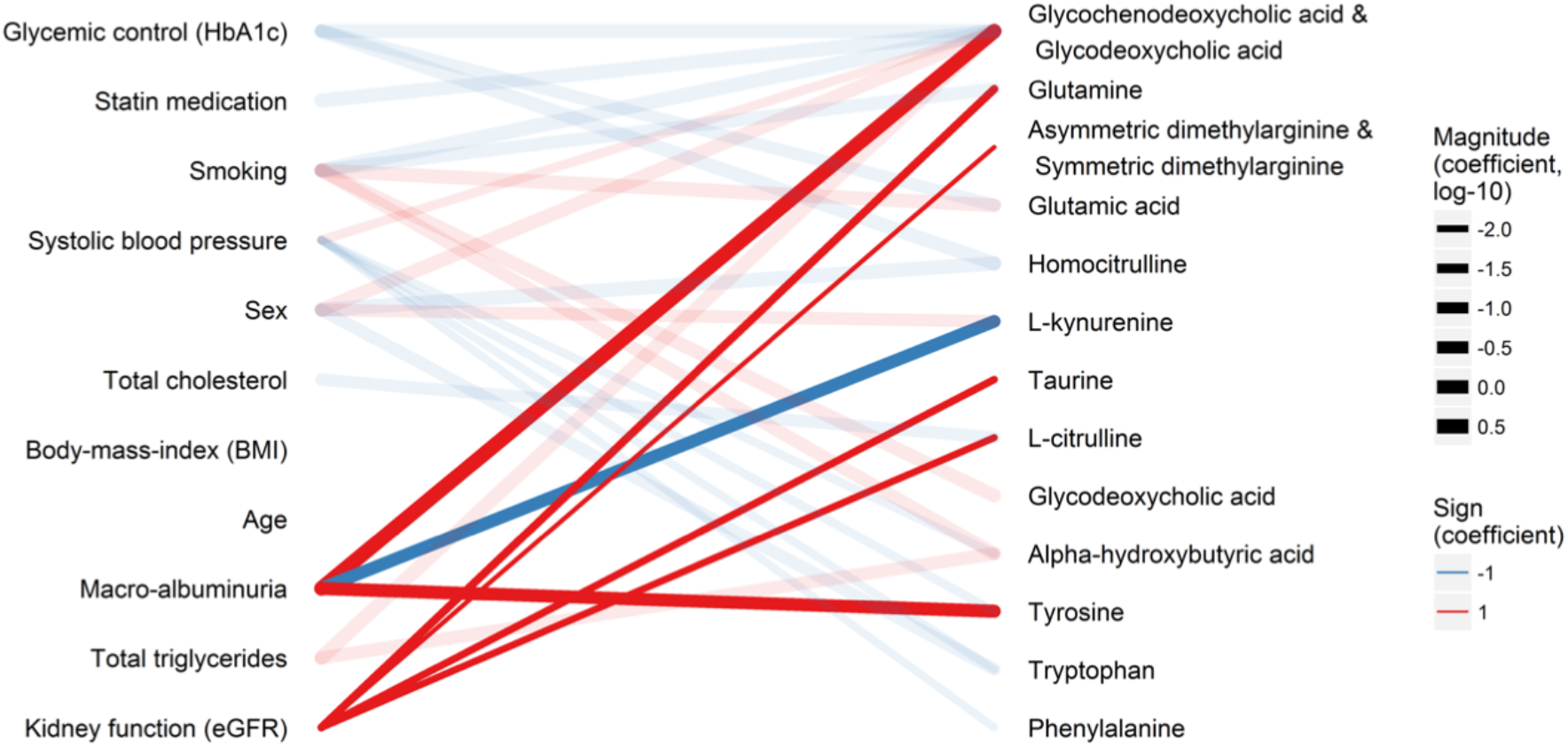
Associations between clinical measurements (left) and the quantified analytes (right) in the T1D cohort. The lines indicate statistical associations (red: positive association and blue: inverse/opposite association; line width: strength of the association). Associations directly related to diabetic kidney disease are highlighted with opaque lines.

The target panel included metabolites which have previously been associated particularly with kidney functions The analysis resulted in estimates of concentrations of the measured metabolites in 50 participants with T1D. Macro-albuminuria, which is an indicator of kidney disease, was associated with elevated GCDCA & GDCA, Tyr and decreased Kynu (**Figure 3**). Estimated globular filtration rate (eGFR; kidney function), was associated with ADMA & SDMA, Cit, Gln, taurine and Tyr. Glycated hemoglobin (HbA1c; glucose control) was associated with decreased GCDCA & GCDA, Glu and HCit. Smoking was associated with elevated Glu, α-OHB and decreased Gln as well as to a disruption in the balance of the bile acids GCDCA and GDCA. Although no metabolomic associations were found with age or BMI in this small sub-study, several metabolites were associated with sex, statin medication, systolic blood pressure, total cholesterol and total triglycerides. It should, however, be noted that as our target panel is based on reported markers of (pre)diabetes and diabetic complications, and does not cover the entire metabolome, a comprehensive pathway analysis could be biased and not fully reliable. The quantitative results are presented in **Supplemental Table S3**.

Overall, our results agree with the previously-published results. Elevated levels of phenylalanine and of arginine, citrulline and ornithine have been reported in T2D patients with macroalbuminuria and microalbuminuric patients, respectively *(38).* Additionally, increased levels of plasma homocysteine have been found to be related to macroalbuminuria in diabetic patients *(39).* ADMA, on the other hand, has been suggested as a candidate biomarker for diabetic kidney complications, whilst elevated levels of ADMA have been shown to predict a more accelerated course of renal function loss and promoted the development of renal damage *(14, 15, 40).* Our results are also in line with literature data showing that bile acid metabolism is altered in T2D patients *(41).* Overall, our results demonstrate that the developed analytical method is feasible for performing targeted metabolomic analysis of plasma samples from diabetic patients, and that it can be used for the stratification of diabetic patients.

## Conclusions

The method developed here proved to be fast (with a sample analysis time of less than 10 minutes) and robust, and thus suitable for routine analyses in the diabetes clinic. Validation of the method showed that the selected panel of markers can be effectively used for classification of subjects with diabetic complications, such as macro-albuminuria. Further evaluation of the clinical relevance of the method is now merited in order to evaluate the full potential of this diagnostic panel in the stratification of prediabetes, metabolic and diabetic complications.

## Acknowledgements

We thank to Nina Christiansen and Birgitte Nergaard Roberts for assistance with sample preparation and to Niina Kärkkäinen for help in the laboratory. We acknowledge Lars Ove Dragsted for valuable input during the final validation of the method. We thank Aidan McGlinchey for language editing. This work was funded by the Novo Nordisk Foundation grant NNF14OC0013659 PROTON.

## Author contributions

1. Linda Ahonen: method development and acquisition of data, analysis and interpretation of data, writing the manuscript
2. Sirkku Jäntti: design of the analytical work, method development and acquisition of data, critical revision of the manuscript
3. Claudia Risz: acquisition of data, critical revision of the manuscript
4. Tommi Suvitaival: design of the clinical sub-cohort, analysis and interpretation of data, critical revision of the manuscript
5. Simone Theilade: design and establishment of the clinical cohort, critical revision of the manuscript
6. Peter Rossing: design of the clinical cohort, critical revision of the manuscript
7. Risto Kostiainen: design of the analytical work, critical revision of the manuscript
8. Matej Oresic: design of the work, critical revision of the manuscript
9. Tuulia Hyotyl<äinen: design of the work, development of the methodology, interpretation of data, writing the manuscript

