## Supplemental Material for "Targeted Clinical Metabolomics Platform for the Stratification of Diabetic Patients"

Linda Ahonen et al.

### *Supplemental Material*

**Supplemental Methods**

*1.1. Chemicals*

LC-MS grade water ($H_2O$), methanol (MeOH), isopropanol (IPA), and acetonitrile (ACN) were purchased from Honeywell International Inc. (Morristown, NJ, USA). HPLC grade dichloromethane (DCM), anhydrous ACN, analytical grade formic acid (HCOOH) and reagent grade potassium carbonate ($K_2CO_3$), potassium bicarbonate ($KHCO_3$), sodium hydroxide (NaOH), hydrochloric acid (HCl) and 5-sulphosalisylic acid dehydrate (SSA) were purchased from Sigma-Aldrich (Steinheim, Germany). 6-aminoquinoline-N-hydroxy-succinimidyl carbamate (AQC) for derivatization of amino acids was purchased from Santa Cruz Biotechnology, Inc. (Dallas, TX, USA).

*1.2. Stock solutions and calibration curves*

Stock solutions (4.0 mg $mL^{-1}$) of the analytes and internal standards were prepared by dissolving in 0.1 M HCl, $H_2O$, $H_2O$:MeOH (90:10, v/v) or in MeOH (**Table 1** and **Table 2** and further diluting them with 0.6 M carbonate buffer (pH 8.9) and 1 M NaOH (3:1, v/v)to the following concentration levels: 2.5, 5.0, 7.5, 10.0, 25, 50, 75, 100, 250, 500, 750, 1000, 2500, 5000, 7500, 10000, 25000, 50000 and 75000 ng $mL^{-1}$. 20 µL of an internal standard solution (ISTD MIX) containing each of the internal standards (**Table 2**) was added to all samples. The samples were vortex mixed and 20 µL of a 5 mg $mL^{-1}$ AQC-reagent in anhydrous ACN (at 55 °C) was added for derivatization of the amino acids and related metabolites (**Figure 1**). Finally, the samples were vortex mixed and stored at -80 °C until analysis. The calibration curves were constructed using at least five measuring points and linear regression with 1/x weighing. For α(R)-OHB and α(S)-OHB, only 3 measuring points could be used due to the high LOD of these analytes.

*1.3. Sample analyses and data processing*

The UHPLC system was 1290 Infinity system from Agilent Technologies (Santa Clara, CA, USA) and it was equipped with a multisampler (maintained at 10 °C), a binary solvent manager and a column thermostat (maintained at 40 °C). The multisampler was set to utilize the Multi Wash option as the needle wash. Here two mixtures, ACN:MeOH:IPA:$H_2O$ (1:1:1:1, v/v/v/v) + 0.1% HCOOH and 10% DCM in MeOH, were used for 8 seconds after each injection in order to clean the needle and the needle seat. Finally, the needle and the needle seat were flushed with the initial gradient conditions for 8 seconds. Separations were performed on a Kinetex® F5 column (100 × 2.1 mm, particle size 1.7 µm) from Phenomenex (Torrance, CA,

USA) with a flow rate of 0.4 mL $min^{-1}$ and an injection volume of 2 µL. $H_2O$ + 0.1% HCOOH (A) and ACN:IPA (2:1, v/v) + 0.1% HCOOH (B) were used as the mobile phases for gradient elution. The gradient was as follows: from 0 to 1 min 1% B, from 1 to 1.8 min 1-18% B, from 1.8 to 3.4 min 18-21% B, from 3.4 to 7 min 21-65% B, from 7 to 7.1 min 65-100% B and from 7.1 to 8.9 min 100% B. Each run was followed by a 2.5 min re-equilibration period under initial conditions (1% B).

The mass spectrometer was a 6460 triple quadrupole system from Agilent Technologies. It was interfaced with an Agilent Jet Stream electrospray ionization source. The analytes were ionized in positive or in negative ion mode depending on the properties of the analyte. Nitrogen generated by a Genius 3010 nitrogen generator from PEAK Scientific Instruments Ltd (Inchinnan, Scotland, UK) was used as the nebulizing gas (pressure 29 psi) and as the sheath gas at 250 °C and 6 L $min^{-1}$ and at 310 °C and 9 L $min^{-1}$, respectively. Pure nitrogen (6.0) from Praxair (Fredericia, Denmark) was used as the collision gas. The capillary voltage was set to 3000 V and the nozzle voltage to 1000 V. MS- and MS/MS-spectra (scan range *m/z* 40-600) were acquired for each analyte to select the best precursor and product ions for selected reaction monitoring (SRM) analyses. The fragmentor voltages, collision energies (CE) and cell accelerator voltages were separately optimized for each ion transition of the analytes (**Table S1**, Supplementary material) and the internal standards (**Table S2**, Supplementary material). MassHunter LC/MS Data Acquisition Software (version B.08.02) was used for all data acquisition. For data processing different software were used: MassHunters Quantitative Analysis Software (version B.07.00), Skyline Daily (version 4.1)*(1)* and R*(2)*

Data from the diabetes cohort were processed as follows: (i) peaks were picked in Skyline *(1)*, (ii) resulting peak areas were normalized to matching internal standard peak areas in R, and (iii) the resulting peak area ratios were calibrated to concentrations in R based on metabolite-specific calibration curves run during the analysis sequence.

**Figure S1A.** Structures of compounds of interest, amino acids and amino acid related compounds.

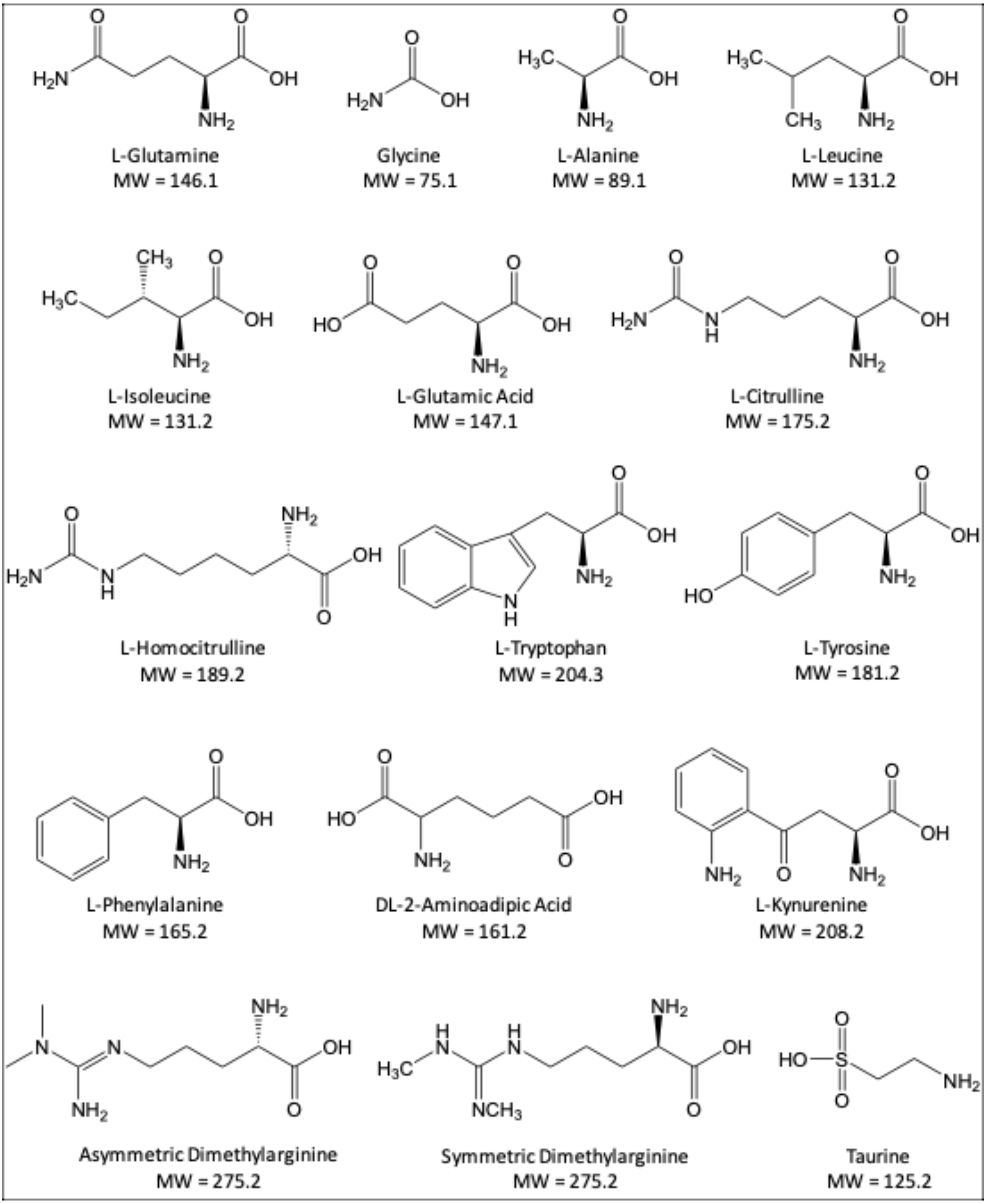

**Figure S1B.** Structures of compounds of interest, bile acids.

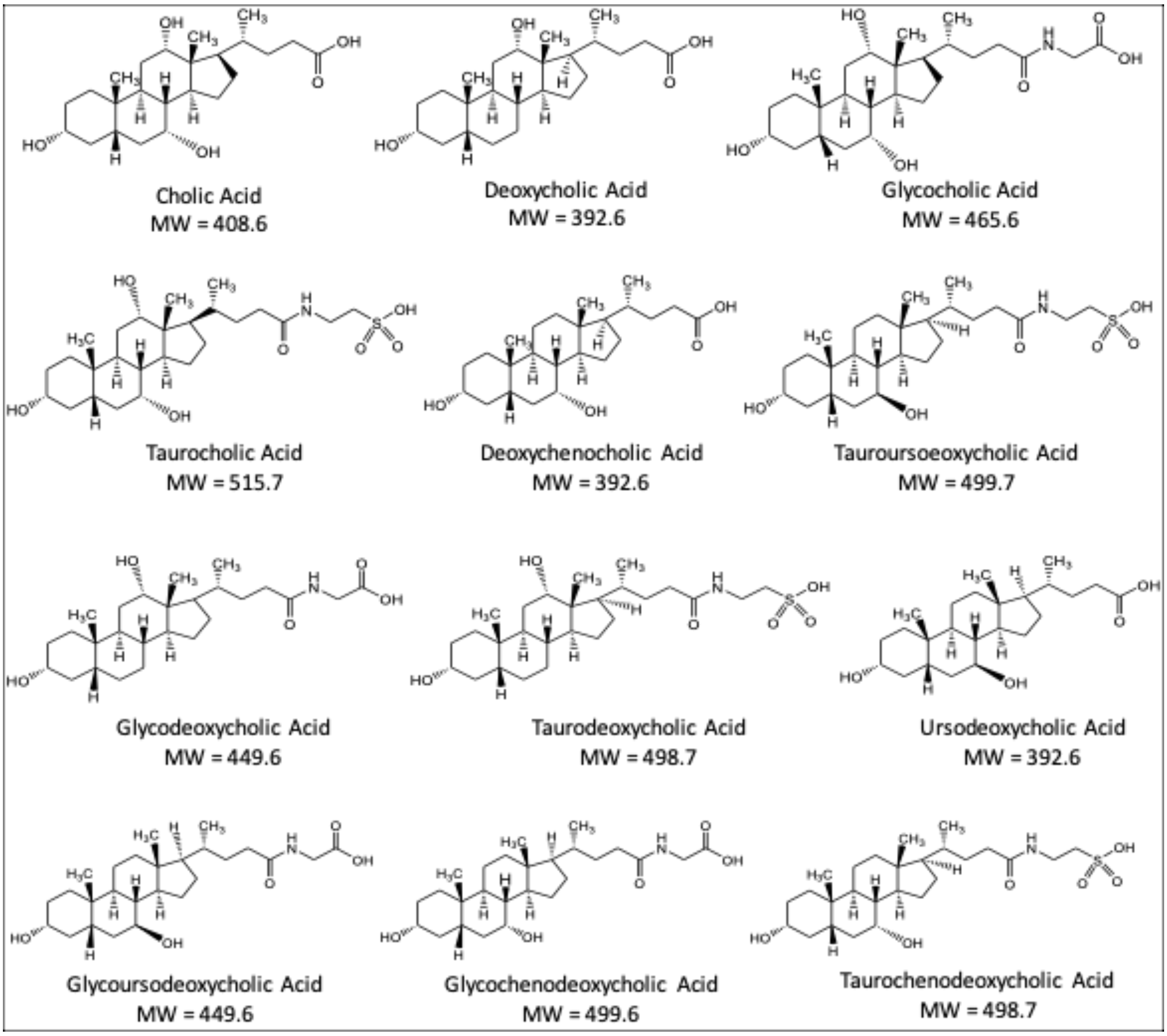

**Figure S1C.** Structures of compounds of interest, small organic acids and other metabolites of interest.

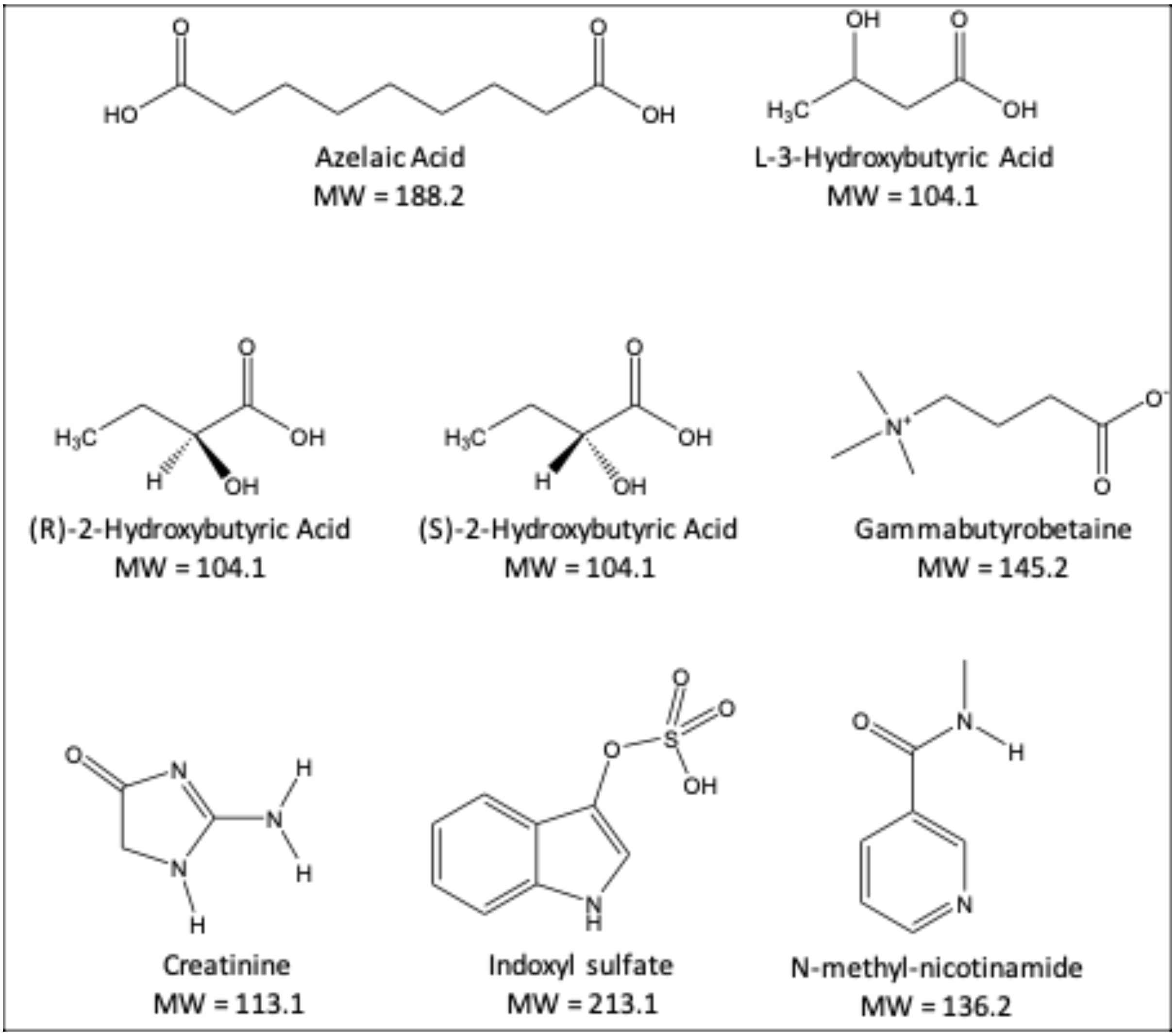

**Figure S2A.** **a)** MS-spectrum and **b)** MS/MS-spectrum of Taurine.

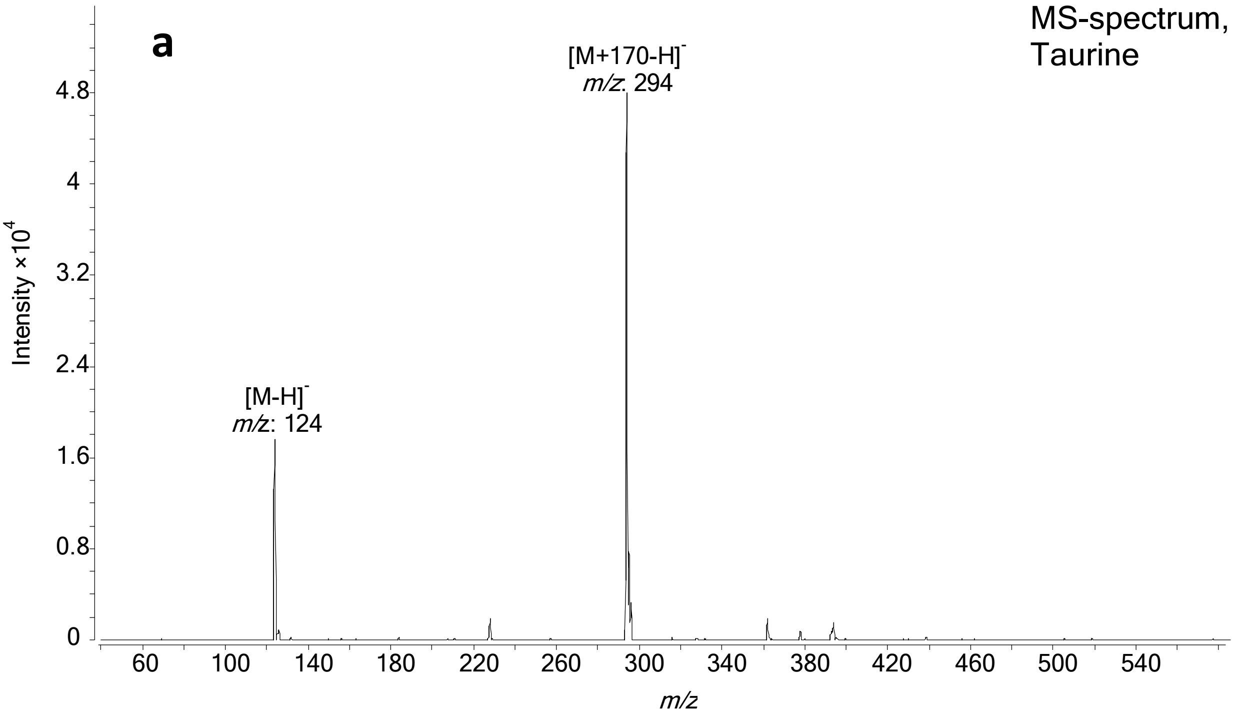


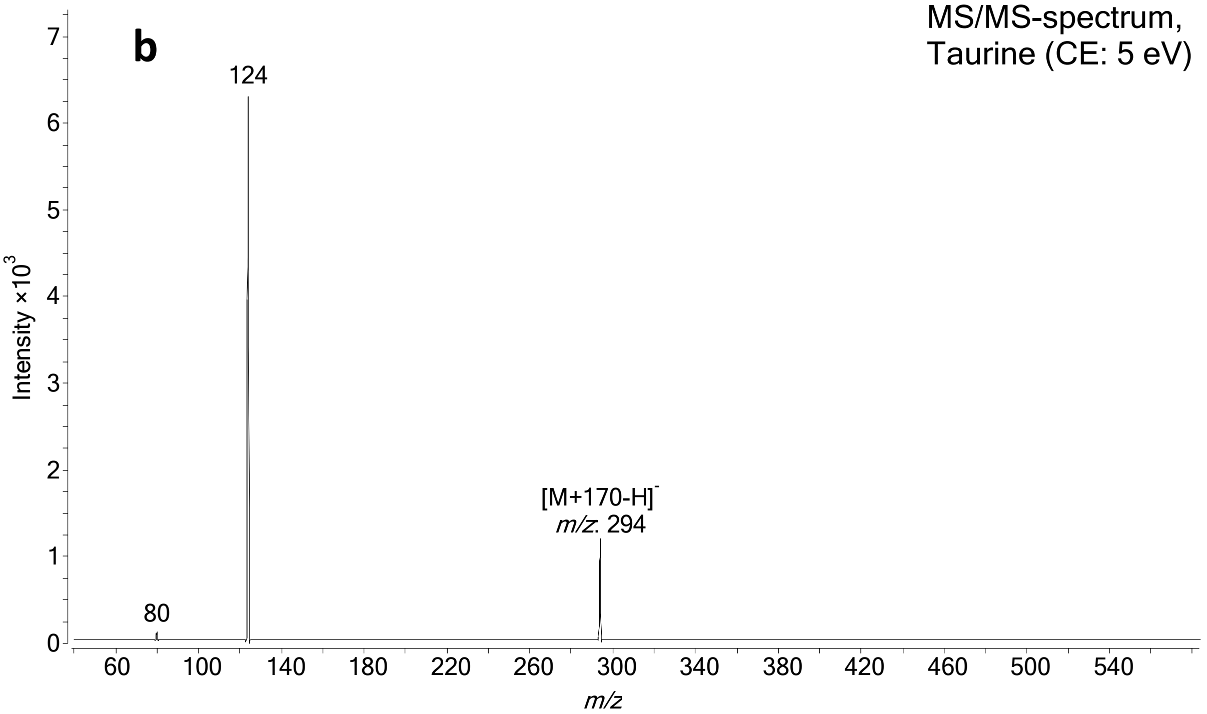

**Figure S2B. a)** MS-spectrum and **b)** MS/MS-spectrum of Azelaic Acid.

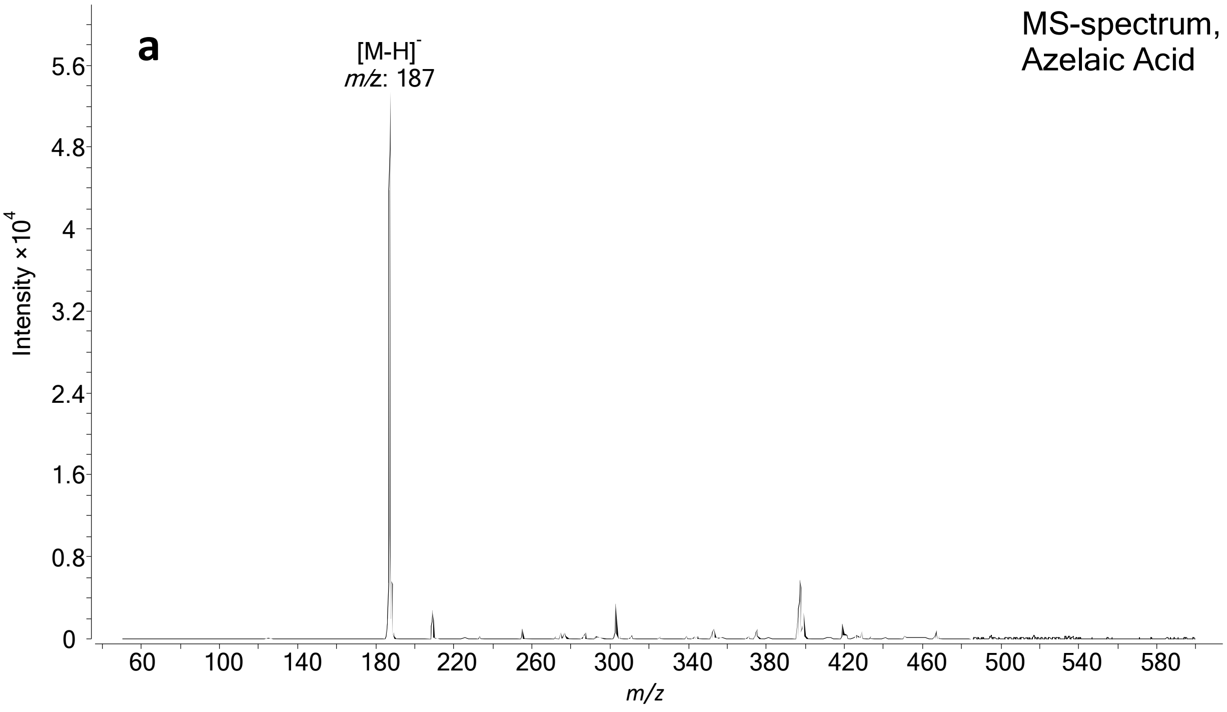


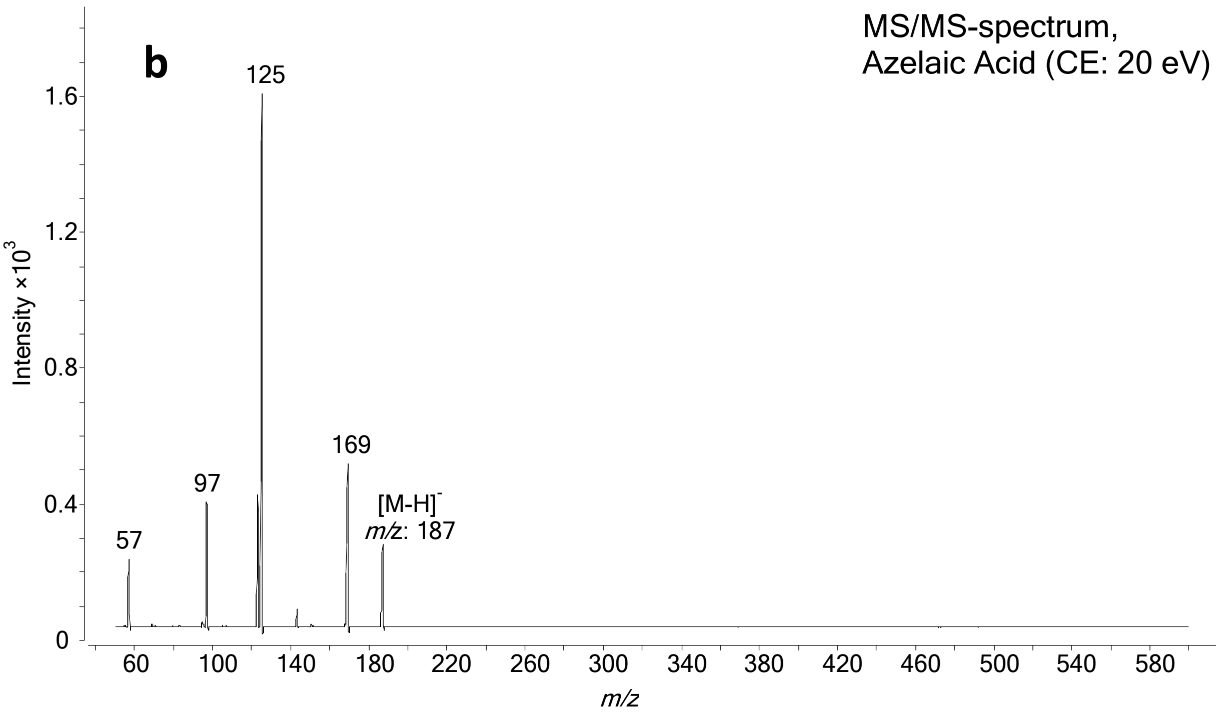

**Figure S2C. a)** MS-spectrum and **b)** MS/MS-spectrum of Gamma-butyrobetaine.

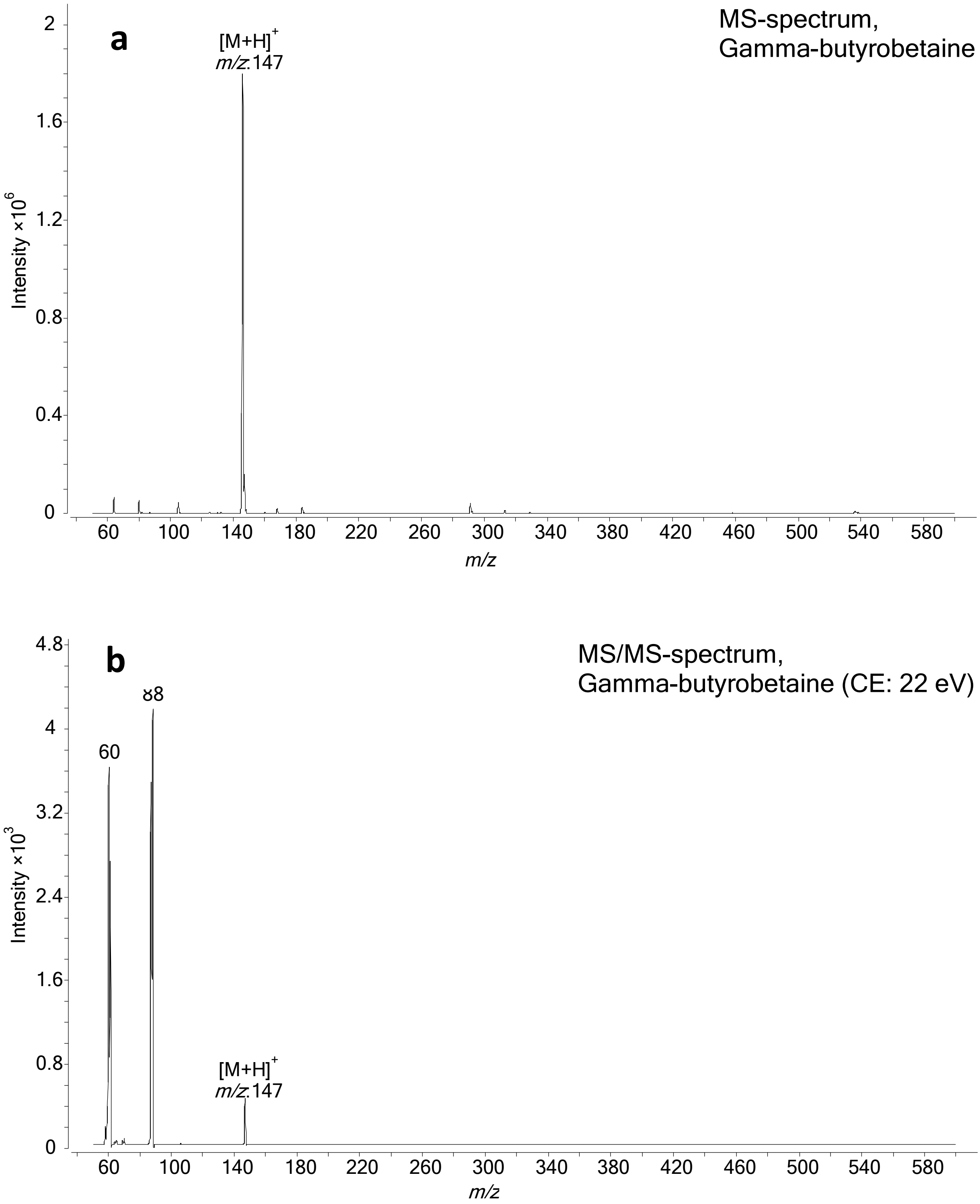

**Figure S2D. a)** MS-spectrum and **b)** MS/MS-spectrum of Glycolic Acid.

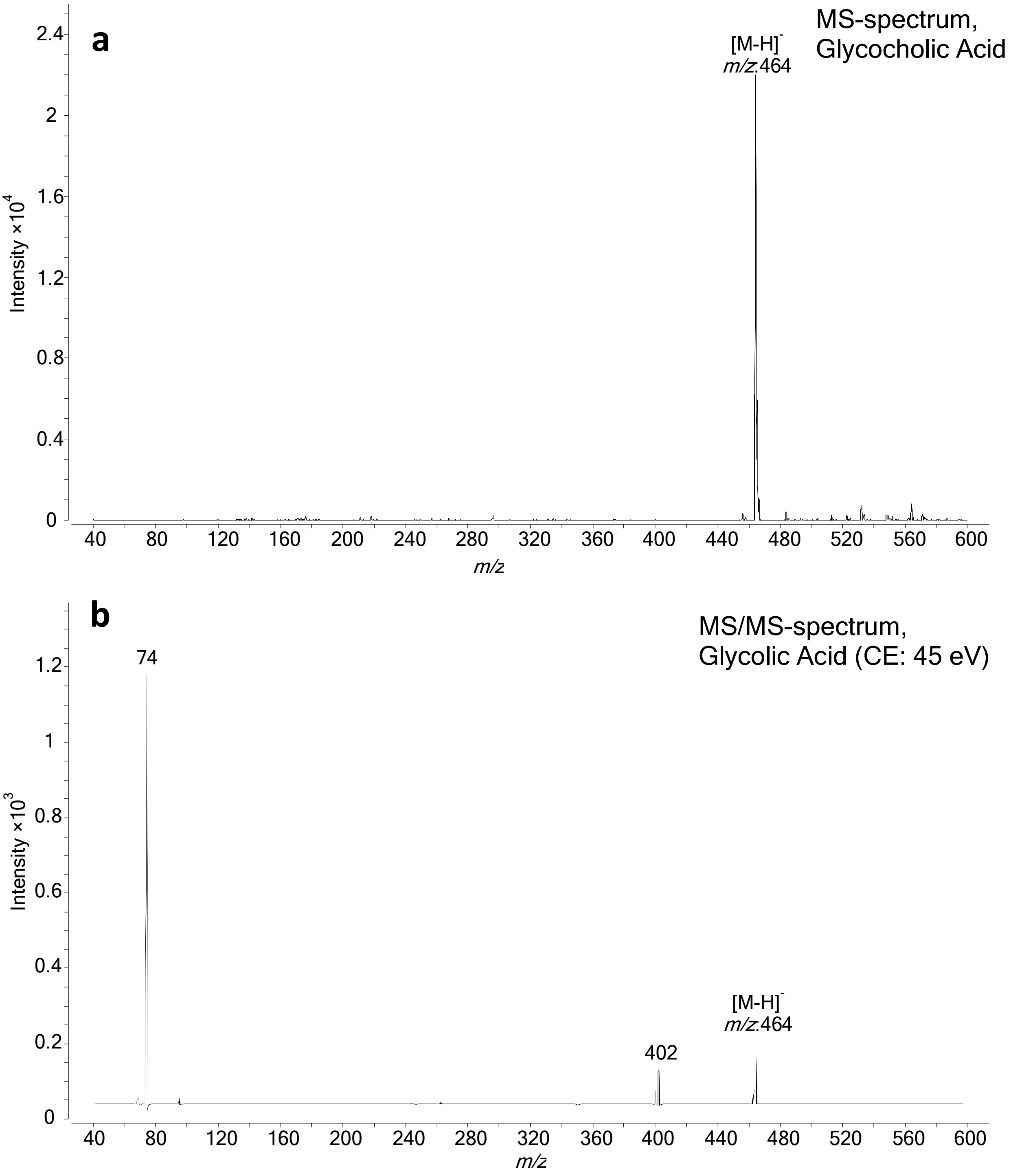

**Figure S2E. a)** MS-spectrum and **b)** MS/MS-spectrum of L-Homocitrulline.

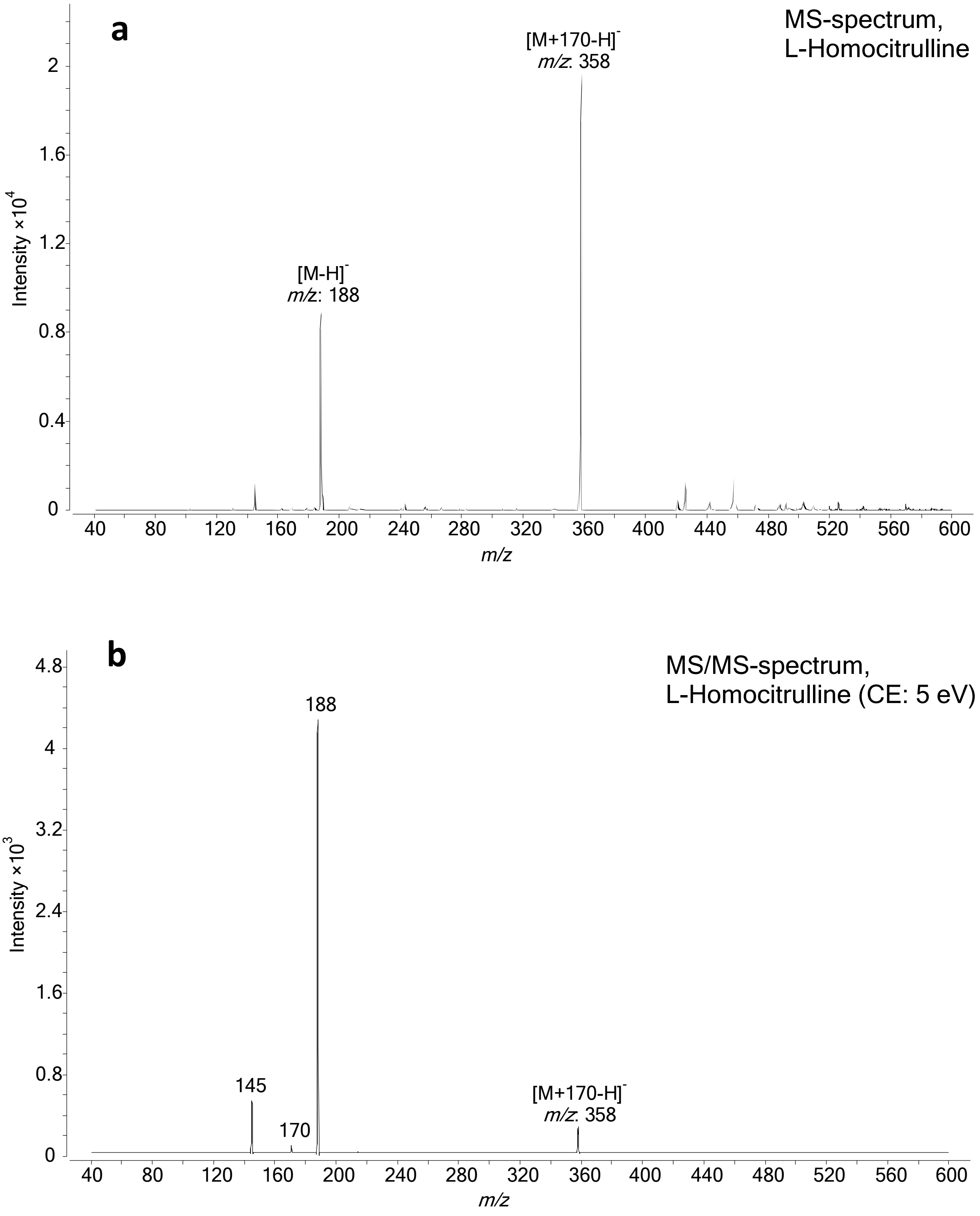

**Table S1.** Optimized fragmentor voltages, collision energies (CE) and cell accelerator voltages for each ion transition of the analytes. The ion transition used for quantification is marked with an *.

| Compound | Molecular weight (MW) | Ion transition | Polarity | Fragmentor voltage (V) | Collision energy (V) | Cell accelerator voltage (V) |
|---|---|---|---|---|---|---|
| **α(R)-OHB & α(S)-OHB** | 104.1 | 103.1 – 45.1 | Negative | 100 | 5 | 4 |
| | | 103.1 – 57.2* | | 100 | 5 | 1 |
| **AADA** | 161.2 | 330.2 – 160.1 | | 150 | 10 | 1 |
| **ADMA & SDMA** | 202.3 | 371.2 – 201.2* | | 150 | 5 | 5 |
| | | 371.2 – 156.1 | | 150 | 20 | 1 |
| **Ala** | 89.1 | 258.1 – 88.1 | | 100 | 15 | 3 |
| **AzelA** | 188.2 | 187.2 – 169 | | 150 | 10 | 1 |
| | | 187.2 – 125.2* | | 150 | 15 | 1 |
| **β-OHB** | 104.1 | 103.2 – 59.2 | | 100 | 5 | 1 |
| **CA** | 408.6 | 407.3 – 407.3* | | 250 | 0 | 1 |
| | | 407.3 – 343.3 | | 250 | 35 | 3 |
| **CDCA** | 392.6 | 391.3 – 391.3 | | 250 | 0 | 3 |
| **Cit** | 175.2 | 344.4 – 174.2 | | 150 | 4 | 7 |
| **Crea** | 113.1 | 114.1 – 86.2 | Positive | 150 | 11 | 4 |
| | | 114.1 – 44.1* | | 150 | 15 | 4 |
| **DCA** | 392.6 | 391.2 – 345.3* | Negative | 200 | 35 | 4 |
| | | 391.2 – 327.2 | | 200 | 40 | 4 |
| **GBB** | 146.2 | 147.2 – 88.1* | Positive | 100 | 16 | 1 |
| | | 147.2 – 60.2 | | 100 | 13 | 1 |
| **GCA** | 465.6 | 464.3 – 402.1 | Negative | 250 | 40 | 4 |
| | | 464.3 – 74.1* | | 250 | 45 | 7 |
| **GCDCA** | 449.6 | 448.3 – 386.3 | | 150 | 40 | 2 |
| **GDCA** | 449.6 | 448.3 – 402.1 | | 250 | 40 | 2 |
| **GCDCA & GDCA** | 449.6 | 448.3 – 74.2 | | 200 | 55 | 2 |
| **Gln** | 146.1 | 315.3 – 145.1 | | 100 | 9 | 6 |
| **Glu** | 147.1 | 316.1 – 146.1 | | 100 | 6 | 6 |
| **Gly** | 75.1 | 244.1 – 74.1 | | 200 | 7 | 4 |
| **GUDCA** | 449.6 | 448.3 – 386 | | 250 | 40 | 2 |
| | | 448.3 – 74.1* | | 250 | 45 | 2 |
| **HCit** | 189.2 | 358.3 – 188.1 | | 200 | 10 | 1 |
| | | 358.3 – 145* | | 150 | 25 | 2 |
| **IndS** | 213.2 | 212 – 132* | | 100 | 15 | 2 |
| | | 212 – 80 | | 100 | 20 | 2 |
| **Kynu** | 208.2 | 377 – 316.1 | | 150 | 5 | 2 |
| | | 377 – 207* | | 150 | 5 | 5 |
| **Leu & Ile** | 131.2 | 300.2 – 130.2 | | 100 | 10 | 1 |
| **N-MNA** | 136.2 | 137.1 – 108.1 | Positive | 100 | 15 | 2 |
| | | 137.1 – 80.2* | | 100 | 26 | 2 |
| **Phe** | 165.2 | 334.2 – 164 | Negative | 100 | 10 | 1 |

| Compound | Molecular weight (MW) | Ion transition | Polarity | Fragmentor voltage (V) | Collision energy (V) | Cell accelerator voltage (V) |
|---|---|---|---|---|---|---|
| **Taurine** | 125.2 | 294.1 – 124.1* | Negative | 100 | 10 | 2 |
| | | 294.1 – 80.1 | | 100 | 55 | 2 |
| **TCA** | 515.7 | 514.3 – 123.8 | | 300 | 65 | 5 |
| | | 514.3 – 80.2* | | 300 | 95 | 1 |
| **TDCA & TCDCA** | 499.3 | 498.3 – 107.1 | | 250 | 80 | 1 |
| | | 498.3 – 80.1* | | 300 | 90 | 1 |
| **Trp** | 204.2 | 373.2 – 203.1 | | 150 | 7 | 2 |
| **TUDCA** | 499.7 | 498.3 – 107.1 | | 300 | 65 | 5 |
| | | 498.3 – 80.1* | | 300 | 85 | 1 |
| **Tyr** | 181.2 | 350.2 – 180.1 | | 100 | 7 | 5 |
| **UDCA** | 392.6 | 391.3 – 391.3 | | 250 | 0 | 4 |

**Table S2.** Optimized fragmentor voltages, collision energies (CE) and cell accelerator voltages for each ion transition of the internal standards.

| Compound | Molecular weight (MW) | Ion transition | Polarity | Fragmentor voltage (V) | Collision energy (V) | Cell accelerator voltage (V) |
|---|---|---|---|---|---|---|
| **AADA-d3** | 164.2 | 333.2 – 145.2 | Negative | 100 | 20 | 2 |
| **ADMA-d7** | 209.8 | 378 – 208.3 | | 100 | 10 | 5 |
| **Ala-d4** | 93.1 | 262.1 – 92.1 | | 100 | 5 | 6 |
| **α-OHB-d3** | 107.1 | 106.1 – 59.1 | | 100 | 10 | 1 |
| **AzelA-d14** | 202.3 | 201.2 – 137.2 | | 150 | 10 | 2 |
| **β-OHB-d4** | 108.1 | 107.1 – 59.1 | | 100 | 5 | 1 |
| **CA-d4** | 412.3 | 411.3 – 411.3 | | 250 | 0 | 3 |
| **CDCA-d4 & DCA-d4** | 396.6 | 395.2 – 395.2 | | 300 | 0 | 4 |
| **Cit-d4** | 179.2 | 348.1 – 135.1 | | 100 | 25 | 2 |
| **Crea-d5** | 118.2 | 119.2 – 49.3 | Positive | 100 | 20 | 1 |
| **GBB-d9** | 154.7 | 155.2 – 87.3 | | 100 | 15 | 6 |
| **GCA-d4** | 469.6 | 468.3 – 74.1 | Negative | 250 | 45 | 1 |
| **GCDCA-d4 & GUDCA-d4** | 453.6 | 452.3 – 74.1 | | 250 | 40 | 1 |
| **GDCA-d6** | 455.7 | 454.3 – 408.2 | | 250 | 55 | 4 |
| **Gln-d5** | 151.2 | 320.1 – 150.1 | | 100 | 5 | 1 |
| **Glu-d5** | 152.1 | 321.1 – 151.1 | | 100 | 5 | 1 |
| **Gly-13C,d2** | 78.1 | 247 – 77.1 | | 100 | 5 | 7 |
| **HCit-2H4** | 193.2 | 362.2 – 192.2 | | 100 | 5 | 6 |
| **IndS-d4** | 217.3 | 216 – 136.1 | | 100 | 15 | 2 |
| **Kynu-13C6** | 214.2 | 383.1 – 195.8 | | 100 | 10 | 6 |
| **Leu-d10 & Ile-d10** | 141.2 | 310.1 – 140 | | 125 | 10 | 2 |
| **N-MNA-d4** | 140.2 | 141.2 – 84.2 | Positive | 100 | 20 | 7 |
| **Phe-d5** | 170.2 | 339.1 – 169.1 | Negative | 150 | 5 | 1 |
| **Taurine-d4** | 129.2 | 298.3 – 128.2 | | 100 | 10 | 3 |
| **TCA-d4** | 519.7 | 518.3 – 80 | | 340 | 100 | 7 |
| **TCDCA-d9** | 508.3 | 507.4 – 80.1 | | 300 | 95 | 1 |
| **Trp-d8** | 212.3 | 381.2 – 211.2 | | 100 | 10 | 5 |
| **TUDCA-d4** | 503.7 | 502.3 – 80.1 | | 300 | 100 | 1 |
| **Tyr-d7** | 188.2 | 357.1 – 187.2 | | 100 | 10 | 1 |
| **UDCA-d4** | 396.6 | 395.3 – 395.3 | | 250 | 0 | 4 |

**Table S3.** Concentrations of metabolites in the validation cohort and their p values.

| Metabolite name | Normo-albuminuria, Mean c (standard deviation) | Macro-albuminuria, Mean c (standard deviation) | P value | adj. p value |
|---|---|---|---|---|
| Glycochenodeoxycholic acid & Glycodeoxycholic acid | 4.33 (11.74) | 2.10 (6.58) | 0.00012 | 0.0021 |
| L-kynurenine | 383.23 (249.28) | 309.03 (86.53) | 0.00043 | 0.0034 |
| Tyrosine | 6185.75 (1865.87) | 7012.51 (2076.69) | 0.00057 | 0.0034 |
| Tryptophan | 5913.04 (1705.38) | 6388.34 (1346.28) | 0.031 | 0.14 |
| Alpha-hydroxybutyric acid | 1464.46 (1276.41) | 1458.64 (1010.47) | 0.055 | 0.17 |
| Glycodeoxycholic acid | 48.77 (6.29) | 51.39 (11.11) | 0.058 | 0.17 |
| Asymmetric dimethylarginine & Symmetric dimethylarginine | 165.73 (51.07) | 153.35 (18.60) | 0.26 | 0.57 |
| Leucine & Isoleucine | 6393.48 (3159.65) | 7303.02 (3656.17) | 0.28 | 0.57 |
| Chenodeoxycholic acid | 1101.07 (7.10) | 1099.58 (6.38) | 0.29 | 0.57 |
| Glycine | 9696.30 (5174.24) | 10313.80 (3604.96) | 0.32 | 0.58 |
| Glutamine | 31651.43 (8920.90) | 29020.85 (6798.27) | 0.4 | 0.63 |
| L-citrulline | 2235.88 (1160.64) | 2253.08 (852.27) | 0.42 | 0.63 |
| Alanine | 16925.72 (4875.55) | 16087.19 (3345.81) | 0.58 | 0.75 |
| Indoxyl sulfate | 907.87 (493.53) | 920.80 (561.30) | 0.6 | 0.75 |
| Homocitrulline | 11.36 (25.76) | 10.21 (20.90) | 0.62 | 0.75 |
| Taurine | 4741.00 (2046.23) | 4128.35 (1424.84) | 0.77 | 0.86 |
| Phenylalanine | 9337.50 (2600.13) | 8949.64 (2062.99) | 0.86 | 0.91 |
| Glutamic acid | 8164.60 (3588.71) | 9304.01 (7562.67) | 0.93 | 0.93 |
